# Metabolite diversity among *Prochlorococcus* strains belonging to divergent ecotypes

**DOI:** 10.1101/2022.12.20.521339

**Authors:** Elizabeth B. Kujawinski, Rogier Braakman, Krista Longnecker, Sallie W. Chisholm, Jamie W. Becker, Keven Dooley, Melissa C. Kido Soule, Gretchen J. Swarr, Kathryn Halloran

## Abstract

The euphotic zone of the surface ocean contains distinct physical-chemical regimes that vary inversely in light and nutrient concentrations as a function of depth. The most numerous phytoplankter of the mid- and low-latitude ocean is the picocyanobacterium *Prochlorococcus,* which consists of ecologically distinct subpopulations (i.e., “ecotypes”). Ecotypes have different temperature, light and nutrient optima and display distinct relative abundances along gradients of these niche dimensions. As a primary producer, *Prochlorococcus* fixes and releases organic carbon to neighboring microbes as part of the microbial loop. However, little is known about the specific molecules *Prochlorococcus* accumulates and releases or how these processes vary among its ecotypes. Here we characterize metabolite diversity of *Prochlorococcus* by profiling three ecologically-distinct cultured strains: MIT9301, representing a high-light adapted ecotype dominating shallow tropical and sub-tropical waters, MIT0801, representing a low-light adapted ecotype found throughout the euphotic zone and MIT9313, representing a low-light adapted ecotype relatively most abundant at the base of the euphotic zone. In both intracellular and extracellular metabolite profiles, we observe striking differences across strains in the accumulation and release of molecules. Some differences reflect variable genome content across the strains, while others likely reflect variable regulation of genetically-conserved pathways. In the extracellular profiles, we identify molecules that may serve as currencies in *Prochlorococcus’* interactions with neighboring microbes and therefore merit further investigation.

**Importance:** Approximately half of the annual carbon fixation on Earth occurs in the surface ocean through the photosynthetic activities of phytoplankton such as the ubiquitous picocyanobacterium *Prochlorococcus.* Ecologically-distinct subpopulations of *Prochlorococcus* (or ecotypes) are central conduits of organic substrates into the ocean microbiome, thus playing important roles in surface ocean production. By measuring the chemical profile of three cultured ecotype strains, we observed striking differences in the likely chemical impact of *Prochlorococcus* subpopulations on their surroundings. Subpopulations differ along gradients of temperature, light and nutrient concentrations, suggesting that these chemical differences could affect carbon cycling in different ocean strata and should be considered in models of *Prochlorococcus* physiology and marine carbon dynamics.

## Introduction

*Prochlorococcus* is the most numerous cyanobacteria species on the planet and fixes ca. 4 Gt of carbon annually (1), or approximately 10% of oceanic carbon fixation. Primary producers such as *Prochlorococcus* anchor the surface ocean microbiome, a consortium of bacteria, phytoplankton, archaea, viruses, and grazers that fix and recycle carbon. Interactions within the microbiome rely on the chemical exchange of metabolites among species, facilitating the turnover of 25 Gt of carbon per year (2). Despite their centrality in the carbon cycle, the identities of most of these metabolites remain elusive, with few constraints on their sources (e.g., *Prochlorococcus*), sinks (e.g., mixotrophic or heterotrophic bacteria), and/or the growth and taxonomic factors that affect metabolite release and composition (3).

One challenge for understanding metabolite production in *Prochlorococcus* is that this genus harbors vast genetic diversity, with unknown impacts on metabolite composition. At the highest level, major branches of the *Prochlorococcus* tree form ecologically coherent populations (“ecotypes”) whose relative abundances shift along large-scale gradients of light, temperature, and nutrients (4-8). A central axis of ecological differentiation is seen most clearly in warm stable water columns: recently-diverging high-light (HL)-adapted ecotypes dominate near the surface, where light is abundant and nutrient levels are low or undetectable, while deeply-branching low-light (LL)-adapted ecotypes increase in abundance deeper in the water column where they encounter low light and elevated nutrients (9-11). Additional niche differentiation *Prochlorococcus* arises due to variations in temperature as a function of latitude (7) and variations in nutrient stresses in different ocean regions (12-14). While the evolutionary diversification of ecotypes has been linked to genetically encoded changes in core metabolic pathways (15), evolutionary changes in other pathways are not yet systematically understood.

These broadly defined ecotypes display significant genomic diversity within coexisting sub-clades. For example, using single-cell genome sequencing, Kashtan et al. (16) showed that two major ecotypes captured within a single drop of water from the North Atlantic contained hundreds of ecologically stable sub-populations that have coexisted for millions of years. More recently, using cyanobacteria-specific amplicon sequencing, Thompson et al. (17) found thousands of variants comprising sub-ecotype populations of *Prochlorococcus* in the north Pacific Ocean. Due to this microdiversity, *Prochlorococcus* has a very large pangenome (i.e., the collective genome of all cells) despite individual cells harboring highly streamlined genomes that lack canonical horizontal gene transfer mechanisms, and instead appear to exploit novel mobile genetic elements called tycheposons (18). Approximately 1000 genes represent the shared core genome (Kettler et al., 2007), with the pangenome of the *Prochlorococcus* ‘collective’ projected to be over 80,000 genes (9). The genetic diversity in the pangenome enables differential responses by cells to nutrient stress due to the presence or absence of specific genes, including for phosphorus (19), nitrogen (20, 21), and iron (22, 23) acquisition. In summary, the hierarchical patterns of diversity reflected in its pangenome have allowed *Prochlorococcus* to achieve a global distribution, high abundance, and ecological stability in the face of myriad and variable environmental forces in the euphotic zone across the world’s oceans (9).

*Prochlorococcus* plays an important role in the global carbon cycle through fixation and subsequent release of organic carbon. However, the composition of the released material, particularly labile molecules such as metabolites, has not been studied systematically, nor have different strains representing diverse ecotypes been compared in this regard. Intracellular metabolites participate in biochemical reactions that power the cell, build biomass, and enable responses to stressors in a changing environment, while extracellular metabolites leave the cell through active and passive release of biochemical by-products, signaling molecules and waste materials (reviewed in 3). In practice, intracellular metabolites reflect near-time response and therefore an instantaneous phenotype, while extracellular metabolites reflect a time-integrated metabolic phenotype (at least in batch culture). Large scale genomic differences among strains or organisms under study result in distinctive metabolite fingerprints in both intra- and extracellular metabolite pools (24, 25). Indeed, metagenomic data from microbial consortia have been used to infer metabolite composition differences within consortia members (26-28). At the strain level, previous work with the human microbiome shows differences in intra- and extracellular metabolite profiles (29-31). In short, variations in metabolite concentrations and composition are expected with genomic variability, but they could also arise from distinct responses to stressors and/or differences in gene regulation and output. The degree to which genomic variability within *Prochlorococcus* ecotypes shapes metabolite composition is currently unknown.

As a first step in understanding metabolite production by *Prochlorococcus,* we chose three strains from phylogenetically and physiologically divergent ecotypes for growth under replete conditions. MIT9301 is a member of the recently-diverged HLII ecotype, which dominates the tropics and sub-tropics at the shallowest depths and is the most abundant *Prochlorococcus* ecotype globally (10). MIT0801 is a member of the LLI ecotype, which is often abundant throughout the euphotic zone with maxima at middle depths, typically 50-100m (32). MIT9313 is a member of the deeply-branching LLIV ecotype that increases with depth, typically reaching its highest abundance below 100 m (10). The LLI ecotype can tolerate light-fluctuations associated with periods of deep mixing during the winter months, when they often become relatively abundant (10, 33). Using mass-spectrometry-based metabolomics, we ask (i) how the intra- and extracellular metabolite composition varies among genetically divergent ecotypes; (ii) how closely patterns in intracellular metabolites reflect genetic variability; and (iii) which extracellular metabolites might be candidates for facilitating interactions with other microbes.

## Results and Discussion

Because the close relatives of our chosen strains generally thrive in different environmental regimes in the wild (10), we considered our metabolite profiles in the context of metabolic processes influenced by key variables in those environments. We examined the role of light by growing all three strains at the same light level and by growing two strains at the light level that more closely approximated their realized optimum in the wild. We tested nutrient stress by growing MIT9301 under low phosphorus conditions. Across our six cultures combining treatments and strains, we quantified 35 intracellular compounds, and 18 extracellular compounds (plus four extracellular metabolites that could not be reliably quantified; Figure 1), representing a subset of conserved metabolic processes of interest in adaptations to light and nutrients.

**Figure 1.**
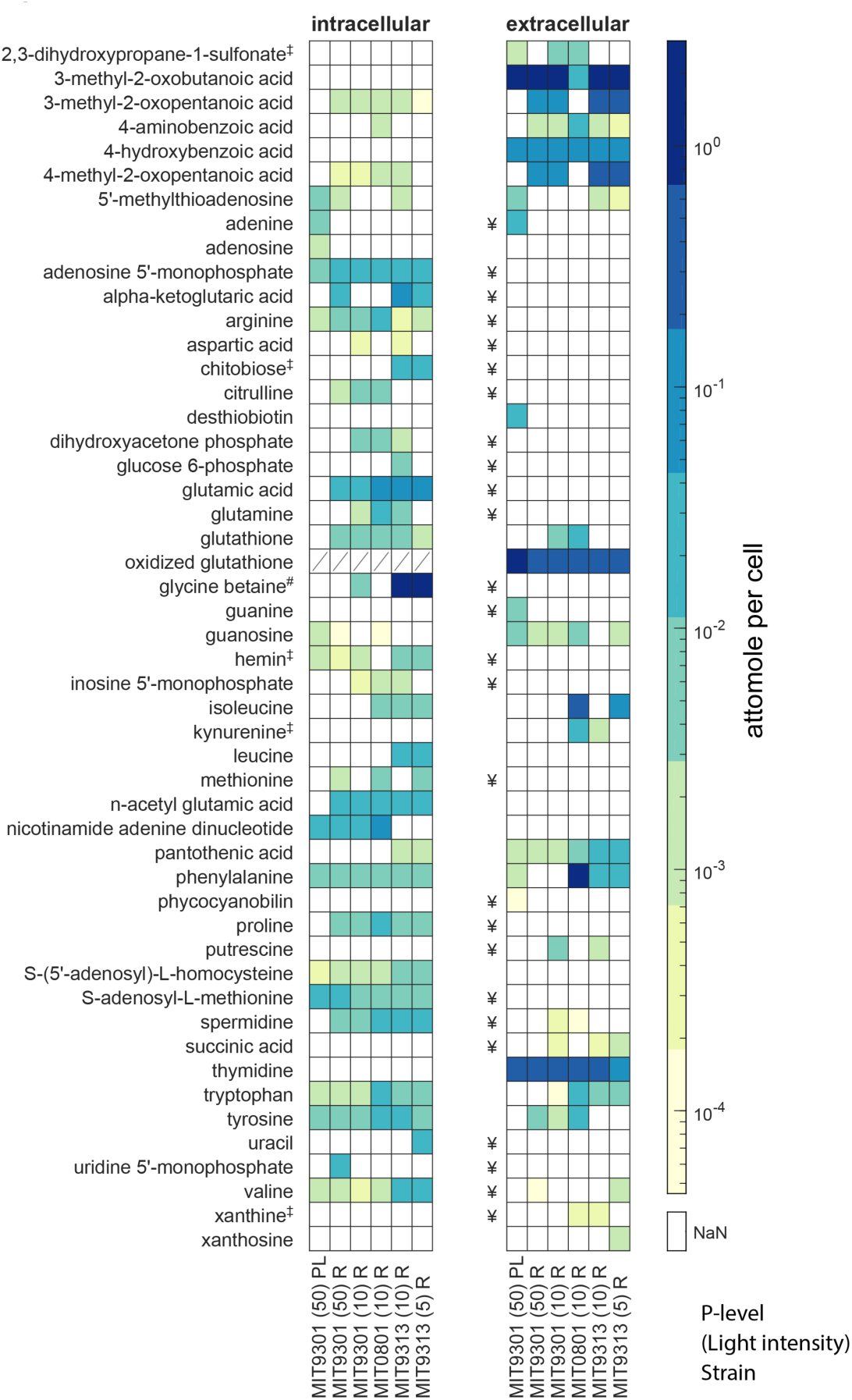
Metabolites detected in three strains of *Prochlorococcus* within intracellular and extracellular phases. Metabolites are presented in alphabetical order with concentrations in attomole per cell; (‡) are metabolites whose biosynthesis genes were not found in the genome, and (#) indicates a metabolite whose biosynthesis genes are present only in MIT9313. Extracellular metabolites with (¥) have an extraction efficiency below 1% and cannot be reliably quantified in the method used here. Intracellular concentrations of oxidized glutathione are not available due to interference by an unknown compound. Columns refer to: MIT9301 (HLII) light intensity 50 µmol photons m^-2^ s^-1^, MIT9301 light intensity 10 µmol photons m^-2^ s^-1^, MIT0801 (LLI) light intensity 10 µmol photons m^-2^ s^-1^, MIT9313 (LLIV) light intensity 10 µmol photons m^-2^ s^-1^, and MIT9313 light intensity 5 µmol photons m^-2^ s^-1^. PL are the phosphate-limited cultures, R is phosphate-replete.

### Metabolite changes linked to differences in light levels and nutrient concentrations

A key parameter that varies among the strains (and ecotypes) is the optimal light intensity for growth, and irradiances below which they cannot survive. MIT9301 (a HLII strain) is better adapted to high light in the surface ocean, displaying optimal growth rates at light intensities that are inhibitory to low-light adapted strains. MIT0801 (LLI) and MIT9313 (LLIV), on the other hand, can survive at light levels that will not sustain high-light adapted ecotypes, and have optimal growth rates at light intensities that are limiting for high-light adapted strains. We explicitly considered light exposure in our experimental design, by growing all strains at an identical intermediate light level and by growing two strains at light levels closer to those where populations in the wild reach their highest densities (Table 1). In each of these two strains, we see few differences in metabolite levels in cultures grown at distinct light levels (Figure 1). Out of 35 intracellular metabolites, only 5 had statistically significant differences in MIT9301 across the two light levels (50 µmol photons m^-2^ s^-1^ and 10 µmol photons m^-2^ s^-1^) and only 3 were statistically significantly different (hereafter simply stated as significantly different) in MIT9313 between 10 µmol photons m^-2^ s^-1^ and 5 µmol photons m^-2^ s^-1^. Out of 18 extracellular metabolites, only 1 was significantly different at different light levels in MIT9301 and only 3 showed significant differences with light in MIT9313.

**Table 1.** Light intensities used to cultivate the strains in this study.

| Strains used in this study | Ecotype | Range of light intensities for ecotype's maximum growth rate in culture<sup>a,c</sup><br>($\mu\text{mol photons m}^{-2} \text{ s}^{-1}$) | Light intensities in this study*<br>($\mu\text{mol photons m}^{-2} \text{ s}^{-1}$) | Light intensity at ecotype's maximum population density in field studies<sup>a,b</sup><br>($\mu\text{mol photons m}^{-2} \text{ s}^{-1}$) |
| --- | --- | --- | --- | --- |
| MIT9301 | HLII | 40-150 | 10 and 50 | 25-85 |
| MIT0801 | LLI | 10-50 | 10 | 2-6.5 |
| MIT9313 | LLIV | 10-50 | 5 and,10 | 0.5-1.5 |

<sup>a</sup>To arrive at these optimal light levels in our study, we examined data from previous laboratory studies conducted under diel light:dark conditions typical of what the cells experience in the oceans. We calculated the total photon fluxes experienced during the light period of diel L:D cycles in those studies, and adjusted light intensities so the cells experienced the equivalent photon flux spread over 24 hours.
<sup>b</sup>Estimated (constant light-equivalent) light level for the water column depth at which populations of different ecotypes reach their maximum density in stratified waters during summer months at HOT and BATS, as determined from Malmstrom et al 2010 (10).
<sup>c</sup>Estimated ranges for the light levels at which growth rates of different ecotypes are maximized in laboratory settings. Not all strains used in this study have been individually characterized, so these ranges reflect the qualitative averages across strains of different ecotypes, as determined in Moore et al (5).

Although we see minimal intra-strain differences in metabolite production as a function of light level, we observed clear inter-strain differences in metabolite levels between strains, including those involved in metabolic processes that could be linked to adaptations or acclimations to different light and nutrient growth conditions (Figures 1 and 2). Two such intracellular metabolites are S-adenosyl-methionine (SAM) and its demethylated product S-adenosyl-homocysteine (SAH), often reported as the ratio of SAM to SAH to highlight the reaction that transfers a methyl group from SAM to other biomolecules, generating SAH. This reaction is a major pathway for methylating DNA, RNA and proteins, a process that often increases under stress (34). Intracellular concentrations of SAM and SAH are tightly regulated to control the extent of methylation of nucleic acids, thus many studies have inferred that higher SAM: SAH values reflect increased methylation potential (35, 36). In this study, we calculated SAM:SAH values for all treatments and strains (Figure 2). We asked two questions in interpreting these data: (i) whether different strains maintained different SAM:SAH values under the same growth conditions and (ii) whether increased stress due to light or nutrients affected SAM:SAH values in a subset of strains.

**Figure 2.**
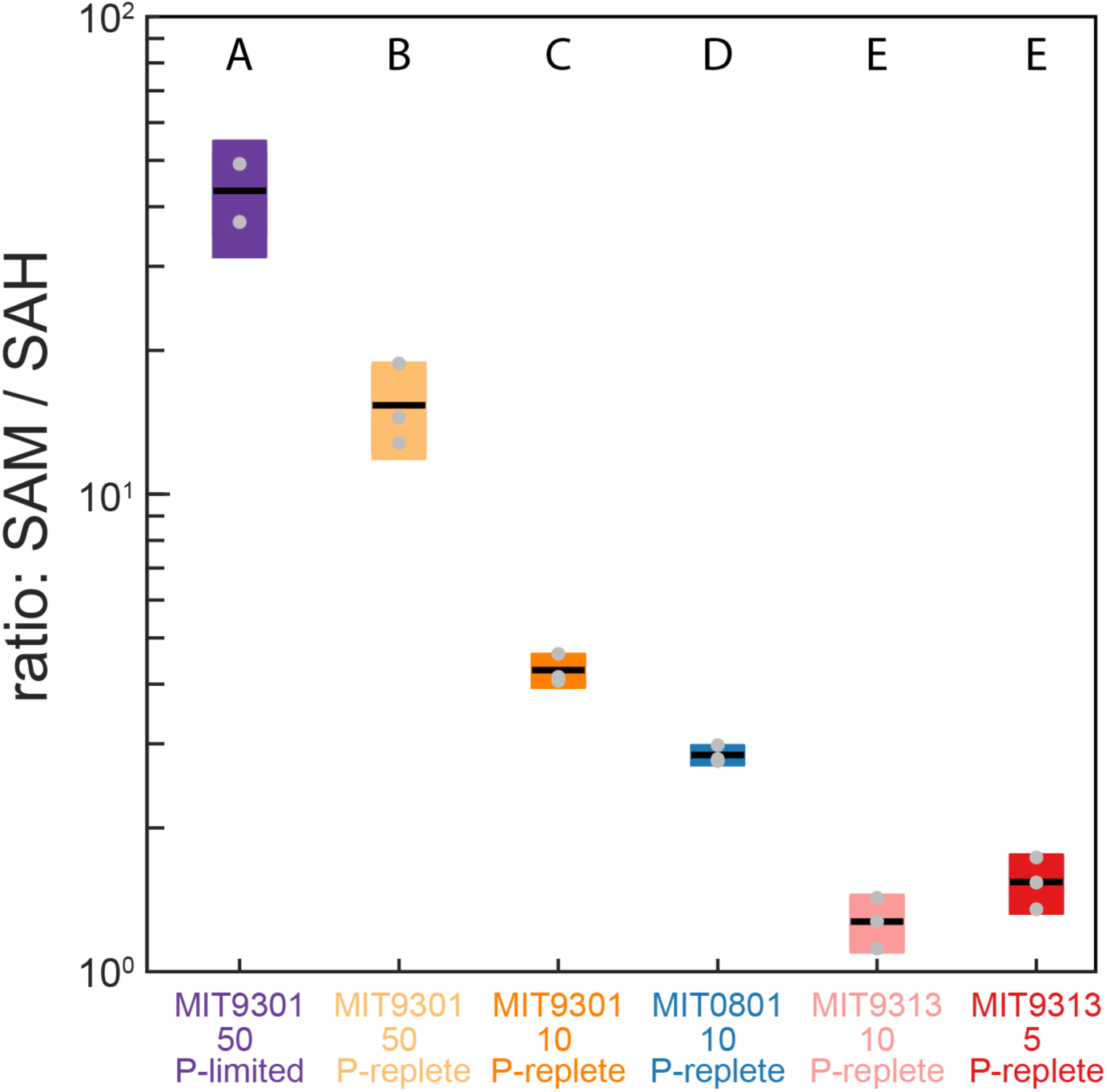
S-adenosyl methionine (SAM) to S-adenosyl-homocysteine (SAH) ratios in three strains of *Prochlorococcus* under different growth conditions. The light intensities used in this study are in Table 1, and are 5, 10, or 50 µmol photons m^-2^ s^-1^. The box-plots represent replicate cultures within each treatment. Letters above each treatment indicate statistically significant differences (two-way ANOVA followed by post-hoc test using Fisher’s least significant difference).

For the first question, the SAM:SAH values across the three strains grown at the same light level (10 µmol photons m^-2^ s^-1^) are statistically significantly higher for MIT9301 than for either MIT0801 or MIT9313 (Figure 2). To ascertain whether there was a genomic basis for this observation, we searched the three strain genomes (in KEGG) for genes involved in SAM metabolism. The three strains share several enzymes (SAM synthetase (*metK*); SAM:tRNA isomerase (*queA*), and SAM decarboxylase (*speD*)), but only MIT9301 encodes the gene for an SAM-dependent cytosine methyltransferase (EC.2.1.1.199). Coe et al. (37) recently showed that within *Prochlorococcus,* only strains from the HLII ecotype methylate cytosine residues in their DNA, although MIT9301 was not tested explicitly. Previous work in *Arabidopsis* showed dynamic and widespread methylation of cytosine residues as a response to different stress types (38). Together these observations suggest that the elevated SAM:SAH values in MIT9301 compared to the other strains could reflect a distinct ability of the HLII ecotype to methylate cytosine residues as a stress response. We tested the impact of stress on SAM:SAH values by comparing this ratio in the three MIT9301 treatments (Figure 2).

Here, we note an increase first with higher light, then a further increase with P-limitation, consistent with sequential increases in stress.

One of the extracellular metabolites that was elevated in MIT9313 under relatively high light was kynurenine, an oxidation product of the aromatic amino acid tryptophan (Figures 1 and 3). Kynurenine was also excreted by MIT0801 (at higher levels than MIT9313). In plant cells, tryptophan residues in photosystem II proteins can be oxidized to kynurenine by reactive oxygen byproducts of photosynthesis (39). During protein remodeling, kynurenine is replaced with tryptophan and excreted or further degraded. The *Prochlorococcus* pangenome does not contain the pathway for kynurenine production or its downstream degradation, so kynurenine excretion in this study (Figures 1 and 3) suggests that this molecule could be an oxidation product of reactive oxygen species (ROS) in the photosystem proteins, similar to hypotheses raised in previous work with *Synechococcus* (40).

**Figure 3.**
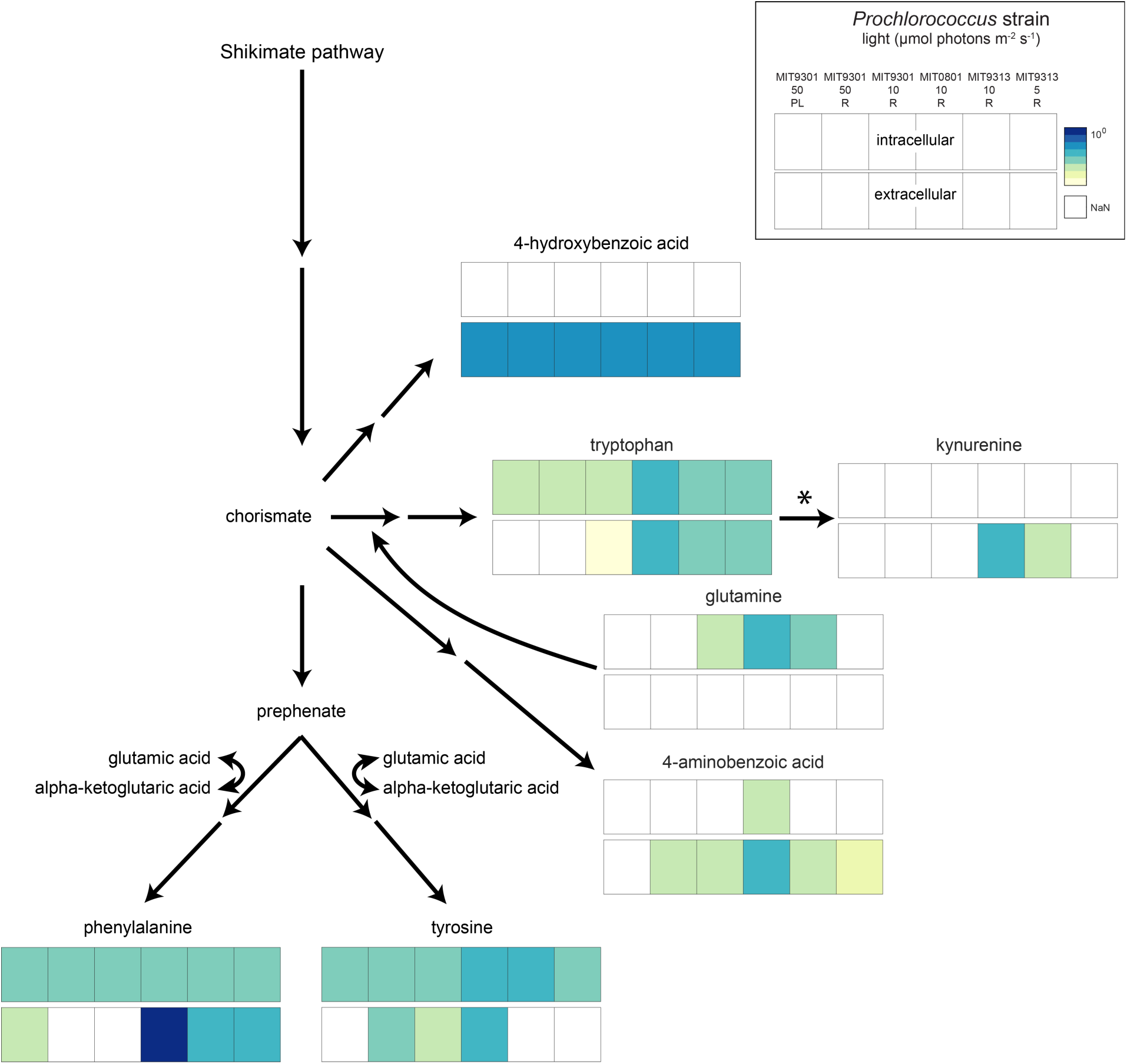
Aromatic amino acids pathway. Squares are log values of mean from Figure 1 (top bar for each compound = Intracellular; bottom bar = Extracellular. Light intensities are shown as well as whether the cultures were P-limited (PL) or replete (R).

As we considered the kynurenine and SAM:SAH dynamics across our study, we asked whether there might be a signal in the aromatic amino acids and their derivatives in the context of *Prochlorococcus* adjusting to different light regimes. The phenyl group is highly reactive to light and reactive oxygen species because the planar π system can adsorb photons and react with oxygen radicals (41, 42). Some microbes, including cyanobacteria, elevate their production of aromatic compounds in response to oxidative stress (43, 44). We observed surprisingly different aromatic amino acid excretion patterns among the strains, despite all three strains having the same relevant biosynthetic pathways (Figure 3, Figure S1). MIT0801 released both tyrosine and phenylalanine, MIT9313 released only phenylalanine and MIT9301 released only tyrosine, indicating possible strain-level idiosyncrasies of these metabolic pathways. However, P-limited MIT9301, for which we inferred enhanced stress from an elevated SAM:SAH value (Figure 2), released phenylalanine instead of tyrosine, suggesting that phenylalanine production and excretion might be a stress response mechanism in *Prochlorococcus* as it is in other systems (43, 44). Further, while levels of tyrosine release were similar for MIT0801 and MIT9301 (∼0.01 amol cell^-1^), levels of phenylalanine release were 30-fold higher in MIT0801 (1.2 amol cell^-1^) than in MIT9313 and P-limited MIT9301, with phenylalanine in MIT0801 being by far the most abundant extracellular aromatic amino acid in our experiment. This raises the possibility that over its unique evolutionary trajectory, MIT0801 has adopted a kind of long-term or constitutively “stressed” state. This speculative conclusion is consistent with the observation that 4-hydroxybenzoic acid, a compound related to aromatic amino acids that is higher in MIT0801 than in the other strains, is also enhanced in MIT9301 under P-limited relative to nutrient replete conditions (Figure 3). However, extracellular oxidized glutathione, a product of peroxide detoxification that exhibits a >10-fold increase in MIT9301 under P-limited relative to nutrient replete conditions, is at similar levels in MIT0801 as in other strains (Figure 1). This suggests that MIT0801 is not actively experiencing enhanced oxidative stress. Regardless of the underlying drivers, differences in observed excretion patterns could be due, at least in part, to differential regulation of the shikimate pathway, which produces aromatic amino acids and is conserved within our three *Prochlorococcus* strains. The first reaction in the pathway is catalyzed by 3-deoxy-7-phosphoheptulonate synthase (DAHP synthetase) and is inhibited by either phenylalanine or tyrosine, leading to excretion of the other (45). How this pathway is regulated in *Prochlorococcus* and under what conditions remains to be determined.

### Metabolite changes linked to changes in nitrogen acquisition and recycling pathways

The ecotypes in our study experience different nutrient concentrations in the wild, with high-light adapted strains typically experiencing lower nitrogen and phosphorus concentrations than the low-light adapted strains. We explicitly tested phosphorus limitation for MIT9301 in this study but kept nitrogen (ammonium) concentrations equivalent among all experiments. We were therefore intrigued to observe higher cell-normalized levels of intracellular amino acids in the LL strains, MIT9313 and MIT0801, than in the HLII strain MIT9301 (Figures 1 and S1). The variability in cell-normalized amino acid concentrations (Figure S1) was surprising given that nearly all amino acid biosynthesis pathways are part of the conserved core genome of *Prochlorococcus* (46). We wondered whether the variability in amino acid levels could reflect the known niche partitioning of *Prochlorococcus* ecotypes along an axis from low nitrogen in the surface ocean (the HL ecotypes) to higher nitrogen deeper in the euphotic zone (the LL ecotypes), specifically in the metabolic processes responsible for nitrogen assimilation, recycling, and storage. We looked first at intracellular citrulline and arginine, two molecules associated with the urea cycle, a common nitrogen recycling strategy (47) and at alpha-ketoglutaric acid and glutamic acid, central metabolites involved in assimilating inorganic nitrogen into protein and nucleic acids (Figure 4). We focused on data from the strains grown under replete conditions to avoid complications associated with phosphorus stress. We observed that citrulline and arginine were positively correlated with one another (R = 0.83; *p* = 0.001), and that alpha-ketoglutaric acid was negatively correlated with citrulline and arginine (R = -0.62 and -0.64, respectively; *p* = 0.03 for both). We observed the highest concentrations of citrulline and arginine in MIT9301 (HLII) and MIT0801 (LLI), and undetectable, or low, concentrations in MIT9313 (LLIV). In contrast, MIT9313 accumulated high concentrations of alpha-ketoglutaric acid and glutamic acid.

**Figure 4.**
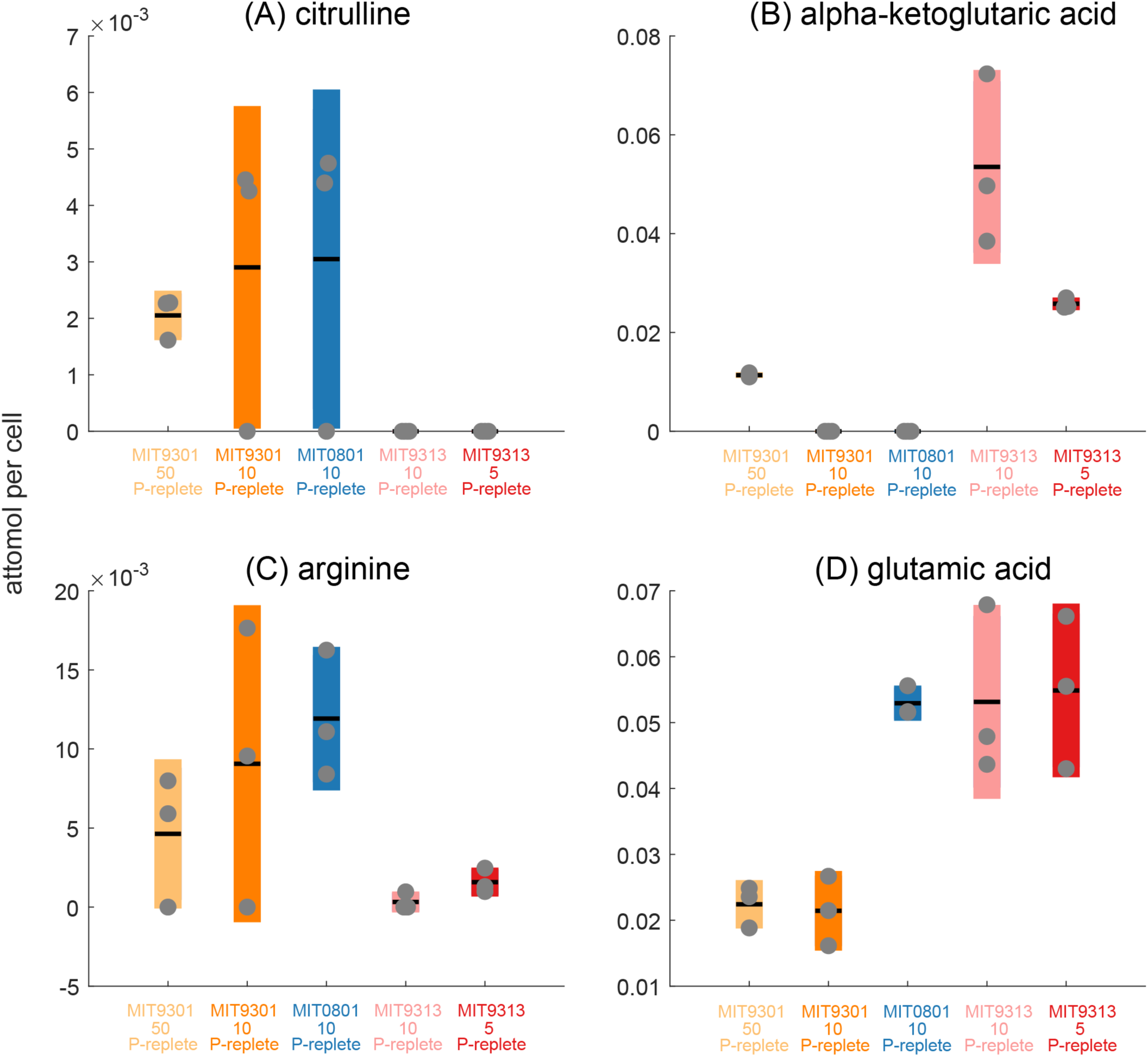
Intracellular cell-specific concentrations of metabolites associated with the urea cycle (A) and (C), compared to metabolites associated with inorganic nitrogen acquisition (B) and (D). Box plots show mean concentrations and standard deviation of experimental triplicates, with grey dots representing individual samples (orange = MIT9301, blue = MIT0801, red = MIT9313).

These amino acids may reflect differences among ecotypes regarding their ability to recycle organic nitrogen and in the properties associated with the two pathways for ammonium assimilation. The increased accumulation of citrulline and arginine, relative to alpha-ketoglutaric acid and glutamic acid, in the high light strain (MIT9301) may indicate a greater tendency to use aspects of the urea cycle by this ecotype, even though *Prochlorococcus* cannot perform the canonical urea cycle due to the absence of the gene for arginase production. Previous field and laboratory work has shown that *Prochlorococcus* uses urea and other forms of organic nitrogen in nutrient-depleted environments, although differences among ecotypes have not been systematically explored (4, 48-51). Further work with nitrogen-limited cultures may provide additional insights into N cycling under these conditions.

To make sense of the alpha-ketoglutaric acid and glutamic acid concentrations in our study, we considered the two pathways for inorganic nitrogen assimilation: (i) the glutamine synthase (GS)-glutamine oxoglutarate aminotransferase (GOGAT) pathway which converts glutamic acid to glutamine and then combines glutamine with alpha-ketoglutaric acid to produce two glutamic acid molecules; and (ii) the glutamic acid dehydrogenase (GDH) pathway, which combines alpha-ketoglutaric acid and ammonium to produce glutamic acid and can work in the reverse direction for organic nitrogen recycling. The GDH pathway is considered a poorer route for glutamic acid biosynthesis because this enzyme has a lower affinity for ammonium, relative to GS. However, the GS-GOGAT pathway consumes ATP while the GDH pathway does not. All *Prochlorococcus* strains encode the high-affinity endergonic GS-GOGAT pathway, but of the strains tested here, only MIT9313 retains the low-affinity but more energetically efficient GDH pathway (46, 52). Flux balance models have shown that low energy-consuming reactions such as the GDH pathway are favored for ecotypes like MIT9313 at the base of the euphotic zone (53).

Both the GS-GOGAT and GDH pathways have been implicated in responses to N stress in transcriptomics, proteomics and metabolomics studies in *Prochlorococcus* (20, 52, 54-56). Taken together, these published studies imply that nitrogen stress induces dynamic responses in transcripts, proteins, and metabolites within the GS-GOGAT pathway for high-light-adapted ecotypes, with more sustained responses in these measurements for low-light ecotypes. For low-light ecotypes, the GDH pathway is not differentially expressed under N stress, and could be important for recycling nitrogen from amino acids (52). In this study, all our strains were grown under replete N conditions, yet we observed elevated alpha-ketoglutaric acid concentrations (Figure 4) in MIT9313 (LLIV), in contrast to MIT9301 (HLII) and MIT0801 (LLI) and previously-published results for VOL29 (HLI; 56). Although transcriptomics data is required to confirm this result, it suggests that the LLIV strain constitutively uses the GDH pathway in addition to the GS-GOGAT pathway to maintain intracellular nitrogen levels, consistent with adaptation of the LLIV clade to an elevated nutrient, energy poor, environment in the deep euphotic zone. Rangel et al. (52) showed that the GDH pathway likely operates in reverse in MIT9313 to transfer ammonium from glutamate, providing a route for utilizing amino acids as a nitrogen source, consistent with findings of elevated mixotrophy in LL-adapted ecotypes (57, 58). Intriguingly, MIT0801 (LLI) also accumulated high intracellular concentrations of glutamic acid, a hint that this strain could be using a combination of nitrogen assimilation strategies (Figure 4).

Another sign that MIT9313 is accustomed to a relatively high-nitrogen environment is its unique ability to synthesize nitrogen-rich organic compatible solutes to offset the inorganic saline ocean. Specifically, we detected very high concentrations of glycine betaine, a common microbial N-containing osmolyte, in MIT9313, consistent with genomic evidence that this strain is uniquely capable of glycine betaine synthesis among our chosen strains and with previous work showing glycine betaine accumulation in MIT9313 (59). In the current study, glycine betaine concentrations reached 3 ± 1.3 fg cell^-1^ or 4-6% of its total biomass (Table S2), supporting its role as an osmolyte in this strain (60). In contrast to MIT9313, *Prochlorococcus* ecotypes living higher in the water column use nitrogen-free osmolytes such as sucrose and glucosyl-glycerate over glycine betaine (60). If glycine betaine production by MIT9313 leads to its excretion, it could promote interactions with neighboring heterotrophs (61), many of which have high-affinity transporters for glycine betaine (62) including the most ubiquitous marine heterotrophic bacteria, SAR11 (63). Indeed, studies of co-cultures of *Prochlorococcus* and SAR11 showed that MIT9313, but not strains from other clades, could fulfill the glycine requirement for SAR11 growth, consistent with excretion of glycine betaine (64).

### Metabolite differences within the branched-chain amino acid (BCAA) pathways

Additional clues to the interplay of stress and metabolite production and excretion come from the branched chain amino acids (BCAAs), valine, leucine and isoleucine and their precursors, which were consistently among the most abundant metabolites measured (Figure 5). When grown under nutrient replete conditions, MIT9301 and MIT9313 excreted high levels of 3-methyl-2-oxobutanoic acid (precursor to valine), 3-methyl-2-oxopentanoic acid (precursor to isoleucine) and 4-methyl-2-oxopentanoic acid (precursor to leucine), but when grown under P-limitation, MIT9301 excreted only 3-methyl-2-oxobutanoic acid. MIT0801, grown under nutrient replete conditions, also excreted only 3-methyl-2-oxobutanoic acid among the three branched chain precursors. These observations suggest *Prochlorococcus* lowers excretion of 3-methyl-2-oxopentanoic acid and 4-methyl-2-oxopentanoic acid under stress but maintains excretion of 3-methyl-2-oxobutanoic acid. Further, they are consistent with the hypothesis that MIT0801 has adopted a kind of constitutively stressed metabolic state. However, as a point of difference in these pathways, in MIT0801 isoleucine was among the most abundant extracellular metabolites, but this compound was undetectable in the other *Prochlorococcus* strains. In general, BCAAs were either not detected or had much lower concentrations than their precursors in MIT9301 and MIT9313, except for isoleucine in MIT9313 grown at low light, where it reached similar levels as its precursor.

**Figure 5.**
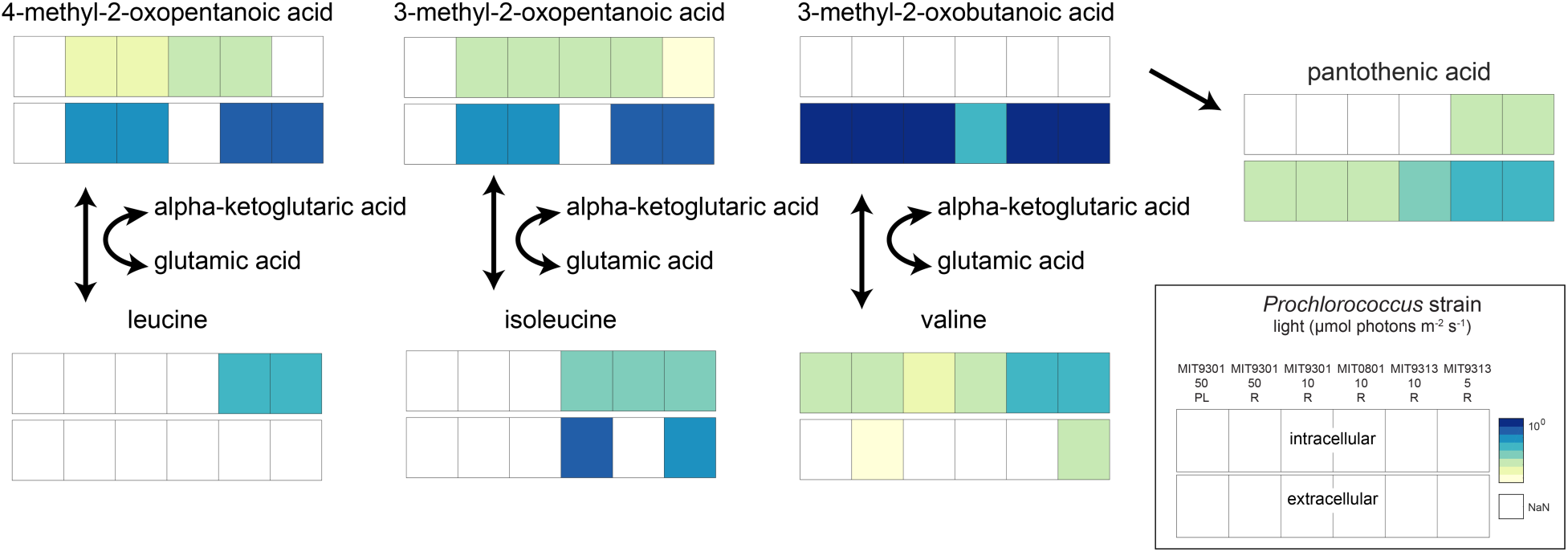
Branched chain amino acids (BCAAs) pathway figure with intermediates and BCAAs. Squares are log values of mean from Figure 1 (top = Intracellular; bottom = Extracellular).

While the above observations suggest that stress modifies the excretion of BCAA precursors, they leave open the question of why *Prochlorococcus* excretes these compounds in the first place. BCAAs play important roles in protein structure due to their hydrophobic and sterically-hindered functionalities and their synthesis pathways are classic examples of allosteric regulation due to the many feedbacks within these pathways (e.g., 65, 66). In all bacteria, BCAA synthesis pathways culminate in the transfer of an amine group from glutamic acid to a BCAA precursor, catalyzed by a shared transaminase. This enzyme also works in the reverse direction and is the first step in BCAA catabolism. *Prochlorococcus* genomes do not contain the full catabolic pathway, but they retain the transaminase step for BCAA amination and de-amination. Consequently, *Prochlorococcus* cannot regulate intracellular BCAA concentrations by catabolizing BCAAs, a common strategy in other microbes (66). Our data cannot distinguish between excretion of these precursors prior to BCAA production, or after the first step of BCAA catabolism. We considered the possibility that excretion of BCAA precursors reflects a form of ‘directed overflow’ metabolism (67), a homeostatic control mechanism in which cells maintain a robust supply of pathway intermediates to ensure efficient synthesis of end-products, and excrete any excess they cannot reuse or recycle. However, in that scenario we might also expect to observe high intracellular levels of the BCAA precursors, but they accumulated at lower levels, if at all, than the BCAAs in all three *Prochlorococcus* strains. The contrast was particularly stark for 3-methyl-2-oxobutanoic acid, which was never detected intracellularly, but its product valine was present in all strains. This may indicate that BCAA intracellular concentrations are regulated to some extent by catabolism to their precursors (to retain intracellular nitrogen), followed by excretion.

Excretion of the nitrogen-free BCAA precursors could therefore reflect large-scale forces shaping *Prochlorococcus* metabolism over the course of its evolution in the nutrient-poor open ocean. That is, reconstructions in the core of carbohydrate metabolism suggest that *Prochlorococcus* has added several pathways for excreting organic carbon as a necessary byproduct of increasing the cellular ATP/ADP ratio to compensate for the increased free energy cost of transporting nutrients from low background concentrations (15). In this context we note that 3-methyl-2-oxobutanoic acid is derived from two molecules of pyruvate, which was identified as one of the central outlets of organic carbon excretion in *Prochlorococcus* (15).

While our observations suggest links between cellular stress and the dynamics of BCAAs, some observed differences across strains are not obviously related to stress, suggesting that other factors and differences between ecotypes are at play. For example, MIT9313 (LLIV) releases the highest levels of 3-methyl-2-oxobutanoic acid and accumulates the highest intracellular concentrations of valine. The justification for high intracellular concentrations of valine and its precursor in the LLIV ecotype, relative to the other strains, is unknown. BCAA precursors could be valuable metabolites to nearby microbes in the field because they require just the transfer of an amino group to generate BCAAs, or because their catabolism can shuttle carbon into core energy-producing pathways. In co-culture experiments with *Prochlorococcus* NATL2A (LLI) and the heterotroph *Alteromonas macleodii*, transcripts for valine catabolism were upregulated in *A. macleodii* relative to a control (68), consistent with cross-feeding of valine and/or its precursor. Although we detected valine in the extracellular phase of two of our strains, this result is treated cautiously due to analytical challenges with the measurement of dissolved valine. Moreover, marine bacteria have not yet been shown to be auxotrophic for BCAAs or their precursors. Nevertheless, our data suggest that these molecules are released at high levels by some *Prochlorococcus* strains and may serve as sources of reduced carbon and/or circumvent the full biosynthetic pathways for BCAAs in nearby heterotrophic bacteria.

When we considered the extracellular dynamics of the BCAA precursors, we noticed that the extracellular concentrations of BCAA precursors were positively correlated with one another and with pantothenic acid (vitamin B5; R = 0.9, *p*-value <<0.001 for pantothenic acid and 3-methyl-2-oxobutanoic acid; Figure 6), while also negatively correlated with thymidine (R = - 0.85, *p*-value <<0.001 for pantothenic acid and thymidine; Figure 6). MIT9313, the LLIV strain, releases higher concentrations of pantothenic acid and BCAA precursors, relative to thymidine. The other two strains have the opposite dynamic, with relatively high thymidine release and lower excretion of pantothenic acid and BCAA precursors. The negative correlation of the three BCAA precursors and pantothenic acid with dissolved thymidine concentrations cannot be explained with any known mechanism. These correlations likely combine differences based on phylogeny with differences in light growth conditions, so it is particularly challenging to identify one explanation. Links between BCAA and thymidine metabolism have been observed in the context of plant herbicide research (69, 70), but these connections require thymidine kinase, an enzyme that is absent in all *Prochlorococcus* genomes. Historically, thymidine was used to assay heterotrophic productivity in the ocean (71), because many microbes can use this compound to make nucleic acids. High levels of thymidine release by *Prochlorococcus* indicate its cross-feeding may be important in the oligotrophic oceans, warranting further study.

**Figure 6.**
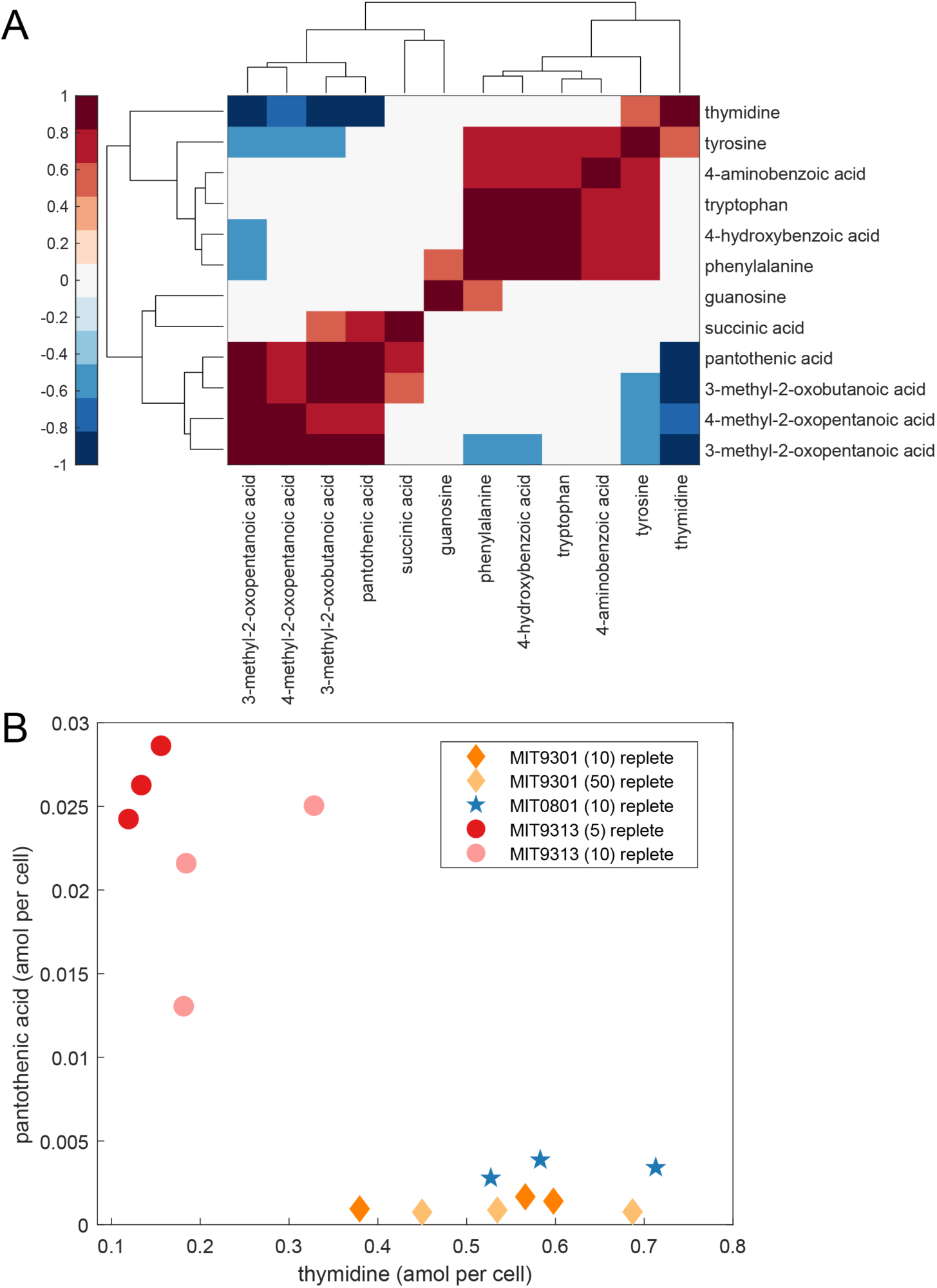
(A) Clustergram of correlations for extracellular (dissolved) metabolites occurring in at least 3 of 5 treatments. The complete set of correlations for dissolved metabolites is provided in Figure S2. Statistically significant positive (red) and negative (blue) correlations are Pearson correlations with p-values adjusted using a False Discovery Rate of 5%. (B) Dissolved pantothenic acid plotted against dissolved thymidine, which are significantly, negatively correlated (orange = MIT9301, blue = MIT0801, red = MIT9313).

Pantothenic acid provides the scaffold for Coenzyme A (CoA), a central metabolite in carbon oxidation cycles, and is one of the most ancient B vitamins due to its ubiquitous role in carbon cycling (72). Further, pantothenic acid is a downstream product of 3-methyl-2-oxobutanoic acid, raising the possibility of a link between the exudation of these compounds, although the underlying mechanism remains unclear. We observed intracellular pantothenic acid only in MIT9313, and as it would likely be quickly converted to CoA, the factors behind its release are not known. However, some ubiquitous marine heterotrophs are auxotrophic for pantothenic acid, including SAR11 and SAR86 (73), suggesting that this molecule may serve an important role in establishing relationships between *Prochlorococcus* and globally abundant sympatric heterotrophic microbes in the surface ocean. Although pantothenic acid has been observed in other culture exudates (74), little is known of its concentrations or dynamics in marine systems (75, 76). The connections observed here among BCAA precursors, pantothenic acid and thymidine hint at potential currencies that could mediate interactions between *Prochlorococcus* and its heterotrophic neighbors.

### Further implications

Stepping back and examining dissolved metabolites in their entirety (Figure S2), we identified additional metabolites that may influence microbial interactions in the ocean. Here, we constrained our interpretation of dissolved metabolites to those whose extraction efficiencies were > 1% (77) and were present in the cultures of at least two strains. In addition to those noted above as present in all strains, five metabolites were detected in the dissolved phase along strain-specific delineations. 5’-Methylthioadenosine (MTA), a byproduct of polyamine metabolism and precursor to methionine salvage, was released by MIT9313 under nutrient replete conditions – albeit at low concentrations – and by MIT9301 under P-limited growth conditions. Release of MTA by MIT9313 is somewhat surprising, as MTA is recycled via the methionine salvage pathway, which is complete only in MIT9313. We hypothesize that *Prochlorococcus* outside of the LLIV clade (including MIT0801 and MIT9301) might instead release 5-methyl-thioribose-1-phosphate (MTRP), a downstream product of MTA that forms a dead end in the network. In this scenario, MTA is released by MIT9301 due to a bottleneck in the availability of phosphate, preventing its conversion to MTRP. MTRP could make for an interesting target in future studies, although it is not currently commercially available and is difficult to synthesize.

We observed release of metabolites whose syntheses are not annotated in any strain’s genome; specifically, 2,3-dihydroxypropane sulfonate (DHPS), xanthine, and kynurenine. All three molecules may be spontaneous reaction products from oxidation or degradation reactions intended for other metabolites. Xanthine and DHPS may be products of nucleic acid degradation (78) and sulfolipid synthesis (79-81), respectively. As noted above, kynurenine is predicted to result from ROS-mediated oxidation of tryptophan (39). Each of these metabolites could be a high-value compound for neighboring heterotrophic bacteria and further work is needed to ascertain whether specific relationships could develop based on these molecular exchanges.

More generally, a notable result in this study is the large variability among metabolite profiles from the three *Prochlorococcus* strains grown under replete conditions. Early metabolomics studies often used one representative strain of model microorganisms for assessing characteristic metabolomes, due to the challenges in collecting and analyzing these data. Indeed, these studies showed that metabolite profiles can reflect differences between organisms with significant genomic variability (e.g., 24). However, our data and previous work (25) suggests that individual metabolite profiles can also be quite different among microbes with more similar genomes. Some metabolite concentrations differed among *Prochlorococcus* strains by a factor of 10X or more in both intracellular and extracellular phases, when normalized by cell number (Figure 1), which suggests differences in regulation, overflow metabolism (67), or other processes that govern how these strains interact with their environment. Some of these differences are larger than those observed in studies within one strain when changing growth conditions (including nutrient-limitation; (56, 82)), underscoring the importance of assessing cross-strain differences in addition to within-strain responses to stressors.

In addition to cross-strain differences, metabolite profiles were generally uncoupled between intracellular and extracellular phases. Direct comparisons between intra- and extracellular phases should be undertaken carefully because the intracellular profiles reflect near-time phenotypes, while the extracellular profiles are the result of accumulation over time in the growth medium. However, if we focus this comparison on metabolites that are equally detectable between the phases (Table S1), we observe cases in which metabolites increase (or decrease) in tandem between the phases (e.g., phenylalanine and pantothenic acid), consistent with internal accumulation driving excretion. But we also see striking differences in the two phases for many metabolites. For example, glutathione and tyrosine are retained with *Prochlorococcus* cells but are detected only in some strains in the extracellular phase. In contrast, thymidine and 3-methyl-2-oxobutanoic acid are not detected intracellularly but are present at high amol cell^-1^ concentrations in the dissolved phase. Thus, *Prochlorococcus* release these metabolites to the external environment without retaining detectable concentrations inside cells. These metabolites could be important currencies in consortia containing *Prochlorococcus*, as they are immediate precursors to nucleic acids, BCAAs, and vitamins. From work with single marine microbial strains, we know that external metabolite profiles are affected by nutrient status (83) and likely by the presence or absence of sympatric heterotrophs (84).

This study represents the first measurements of some of the metabolites we report, limiting our conclusions due to insufficient background information on these compounds’ biochemical roles. Nevertheless, these data suggest that molecules that are transferred among microbes may not be easily predicted from particulate measurements or from genomic considerations. Instead, dissolved metabolite analyses will be needed to establish the dominant currencies of the marine microbial loop, and some of the molecules observed and described here are excellent targets for field, laboratory, and modelling efforts.

## Materials and Methods

### Strain information

The strains in this study included MIT9301 (HLII; 85), MIT0801 (LLI; 32), and MIT9313 (LLIV; 8). Genomes for each of these strains are publicly available and the Kyoto Encyclopedia of Genes and Genomes (KEGG) contains information on their known biochemical pathways: MIT9301 = ‘pmg’ in KEGG, MIT0801 = ‘prm’, and MIT9313 = ‘pmt’.

### Culture conditions

We cultured all three strains individually in 0.2-µm filter-sterilized Sargasso Sea water amended with sterile Pro99 nutrients (86) at 24 °C under constant illumination. We chose light levels to balance maximizing growth rates *in vitro* (5, 8) and conditions relevant to different ecotypes in the wild (10), and to facilitate light-independent comparisons among strains – i.e., when possible growing all three strains at the same light intensity (Table 1). For example, all three strains were grown at 10 µmol photons m^-2^ s^-1^, which lies between the optimal level to achieve maximum *in vitro* growth rates for members of the LLI ecotype (MIT0801) and conditions in the mid-euphotic zone where populations of the LLI ecotype typically thrive. Since the HLII and LLIV ecotypes generally inhabit the upper and lower euphotic zone, respectively (10), we also acclimated MIT9301 to a higher irradiance (50 µmol photons m^-2^ s^-1^) and MIT9313 to a lower irradiance (5 µmol photons m^-2^ s^-1^).

To assess the impact of stress associated with nutrient limitation, we grew MIT9301 at the higher irradiance (50 µmol photons m^-2^ s^-1^) under semi-continuous phosphate limitation through daily transfers (at a dilution rate of 0.3 day^-1^) in media with 20-fold lower phosphate concentrations (i.e., 2.5 µM phosphate vs 50 µM phosphate) than the nutrient-balanced Pro99 media used for the other cultures. We maintained each strain at its target irradiance levels (+/- 5%) for >50 generations in triplicate axenic batch cultures in acid-washed autoclaved borosilicate glass tubes before sampling. We confirmed purity by flow cytometry and a suite of purity test broths (ProAC, ProMM, and MPTB) as previously described (5, 21, 87-90). We monitored growth via bulk fluorescence measurements, and enumerated cells using a Guava Technologies easyCyte 12HT flow cytometer (EMD Millipore) measuring red fluorescence (695/50) and forward scatter from blue (488 nm) excitation. We diluted samples in sterilized Sargasso Sea water prior to enumeration to ensure <500 cells µL^-1^ and avoid coincidence counting.

### Metabolite sampling and extractions

We sampled triplicate batch cultures (35 mL) in mid-exponential growth for both intracellular and extracellular metabolite profiles for all replete treatments. For the P-limited MIT9301 cultures, cell densities were significantly lower due to the phosphate limitation, requiring us to sample over several days – as dictated by the dilution rate – to obtain an approximately similar total number of cells as from the replete cultures. Specifically, we sampled 9 mL (i.e., the volume of the culture replaced with fresh media each day) from the semi-continuous cultures for 6 consecutive days before destructively sampling the full remaining volume. All daily samples from each replicate culture were combined after completion of the experiment. Due to this “averaging” effect of combining the daily samples and the more complex sampling scheme, we opted to obtain duplicates rather than triplicates for the P-limited cultures. All glassware used in metabolite extractions was acid-washed, rinsed with ultrapure water (EMD Millipore), and combusted (450 °C for 8 h) prior to use. We filtered each replicate through a 0.1-µm Omnipore filter (Millipore) in a glass filtration tower under a gentle vacuum and exact volumes were recorded with sterile serological pipettes. We used the filters for intracellular metabolites; they were folded (sample side in), placed in cryovials, and immediately stored at -80 °C. We used the filtrates for extracellular metabolites; they were transferred to acid-washed, autoclaved polycarbonate bottles and acidified with 1 µL of 12 M HCl per mL of sample (final pH ≈ 3) before storage at -20°C.

We extracted intracellular metabolites using a method described by Rabinowitz and Kimball (91) and modified as described in Kido Soule et al. (92). Briefly, we extracted each filter three times with ice-cold extraction solvent (40:40:20 acetonitrile: methanol: water with 0.1 M formic acid). We neutralized the combined extracts with 6M ammonium hydroxide and dried them in a vacufuge until near dryness.

We extracted extracellular metabolites from the seawater matrix using Bond Elut PPL cartridges (1 g/6 ml sized cartridges, Agilent) following the protocol of Dittmar et al. (93) as modified by Longnecker (94). We eluted dissolved metabolites from the cartridges with 100% methanol and stored them at -20 °C until analysis. Immediately prior to analysis, we dried (to near dryness) the extracts with a vacufuge.

### Targeted mass spectrometry

We re-constituted all extracts (intracellular and extracellular) in 95:5 (v/v) water: acetonitrile with isotopically-labeled injection standards (50 pg/µL each; d2 biotin, d6 succinic acid, d4 cholic acid, d7 indole-3-acetic acid and ^13^C1-phenylalanine). To quantify targeted metabolites, we used ultra-performance liquid chromatography (Accela Open Autosampler and Accela 1250 Pump, Thermo Scientific) coupled to a heated electrospray ionization source (H-ESI) and a triple quadrupole mass spectrometer (TSQ Vantage, Thermo Scientific) operated under selected reaction monitoring (SRM) mode. We performed chromatographic separation with a Waters Acquity HSS T3 column (2.1 × 100 mm, 1.8 µm) equipped with a Vanguard pre-column and maintained at 40 °C. We eluted the metabolites from the column with (A) 0.1% formic acid in water and (B) 0.1% formic acid in acetonitrile at a flow rate of 0.5 mL min^-1^, according to the gradient: 0 min, 1% B; 1 min, 1% B; 3 min, 15% B; 6 min, 50% B; 9 min, 95% B; 10 min, 95% B; 10.2 min, 1% B; 12 min, 1% B (total run time = 12 min). For positive and negative modes, we performed separate autosampler injections of 5 µL each.

We separated extracts into intra- and extra-cellular batches and generated a pooled sample using 25 µL of each sample. Within a batch, we analyzed the samples in random order. We operated the mass spectrometer in SRM mode with optimal SRM parameters (s-lens, collision energy) for each target compound determined previously with an authentic standard. The complete list of metabolites is provided in Table S1. We monitored two SRM transitions per compound for quantification and confirmation and generated 10-point external calibration curves based on peak area for each compound. We converted raw data files from proprietary Thermo (.RAW) format to mzML using the msConvert tool (95) prior to processing with MAVEN (96). We required metabolites to be detected in two of three biological replicates (or in both replicates of the P-limited MIT9301 cultures) to be considered for further analysis.

Extracellular metabolite concentrations were corrected for extraction efficiency (77). We report all detected metabolites, but only concentrations for those with extraction efficiencies >1%.

Extracellular (dissolved) and intracellular (particulate) metabolite abundances are presented as cell-specific concentrations, calculated as the moles of each metabolite divided by *Prochlorococcus* cell number.

### Statistics, data, and code availability

We confirmed that the target metabolites in this study could be synthesized by at least one strain using the KEGG module within Biopython (97). Because KEGG is not organized to consider strain-specific differences in metabolites, we queried KEGG to find the genes unique to each strain, then to determine the list of reactions associated with a given gene, and finally, to identify chemical compounds (metabolites) associated with each pathway.

For MIT9301 and MIT9313, *Prochlorococcus* was grown at two light levels and we tested for differences due to light with a t-test where *p*-values were adjusted for multiple comparisons allowing a False Discovery Rate of 5% (98). Correlations between metabolites were calculated as Pearson correlations and *p*-values were adjusted using a False Discovery Rate of 5% (99). A two-way ANOVA using log10-transformed data followed by post-hoc tests using Fisher’s least significant difference were used to consider differences in the ratio of metabolites across ecotype and light level. Clustergrams were calculated in MATLAB 2019b using a Euclidean distance metric and average linkage. Select figures were made using the notBoxPlot function from the MATLAB File Exchange; these figures show discrete datapoints, the mean value, and the 95% confidence interval for the mean.

Targeted metabolomics data for this project are available from MetaboLights (https://www.ebi.ac.uk/metabolights/MTBLS567). The code to query genetic data based on metabolomics information is available at GitHub (https://github.com/KujawinskiLaboratory/Pro_mtabs/blob/master/find_metabolites_by_strain.ipynb)

## Acknowledgements

The mass spectrometry samples were analyzed at the WHOI FT-MS Users’ Facility with instrumentation funded by the National Science Foundation (grant OCE-1058448 to EBK and MCKS). This work was supported in part by grants from the Simons Foundation (Award ID #509034 to EBK, Award ID #509034SCFY20 to RB and SWC, and SCOPE Award ID 030793 to SWC and Award ID 329108 to M.J. Follows).

## Supplemental information

The supplemental information includes 2 tables and 2 figures.

**Figure S1.**
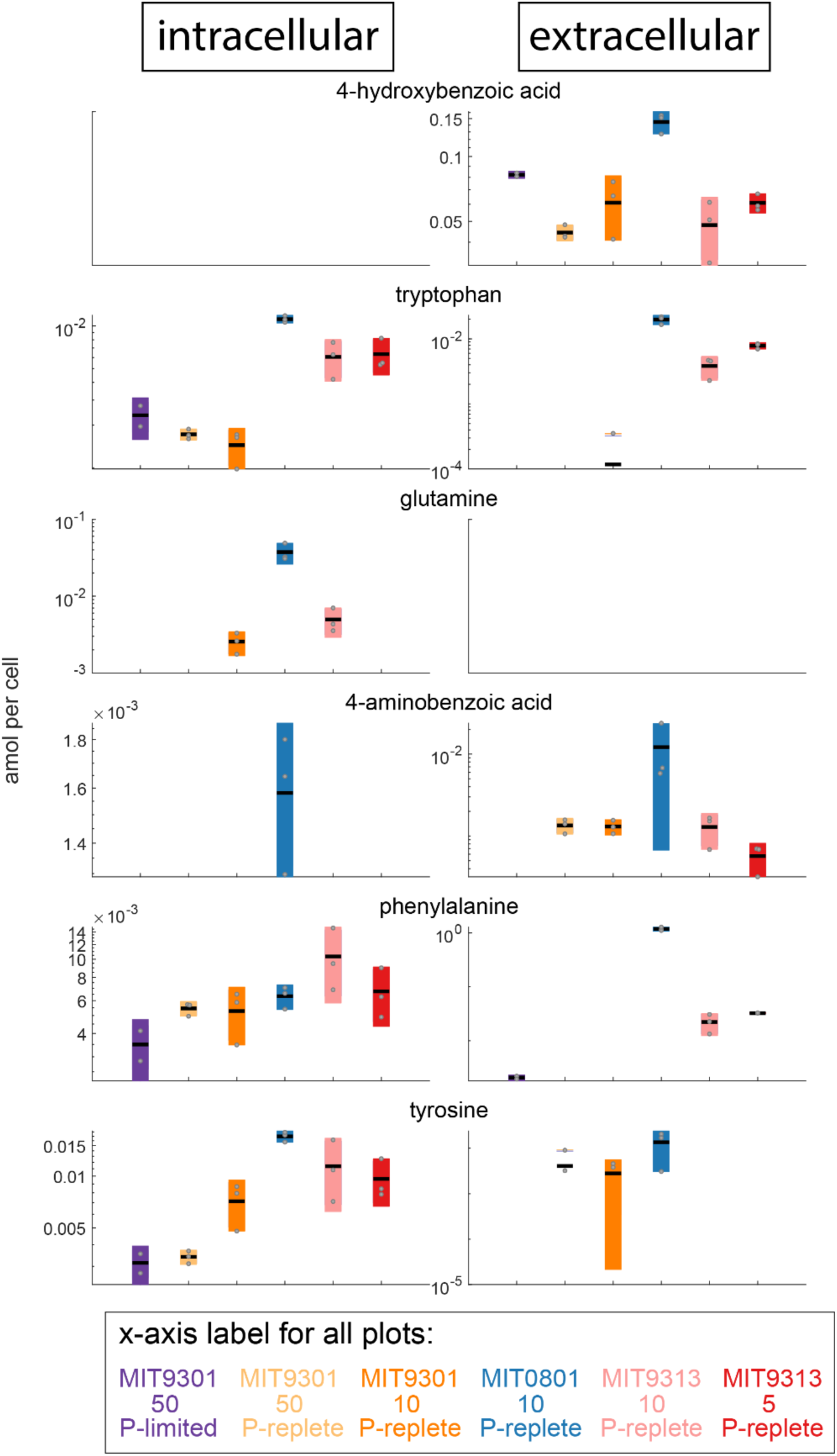
Metabolites from Figure 3, plotted on a log scale as discrete amol per cell values for each metabolite.

**Figure S2.**
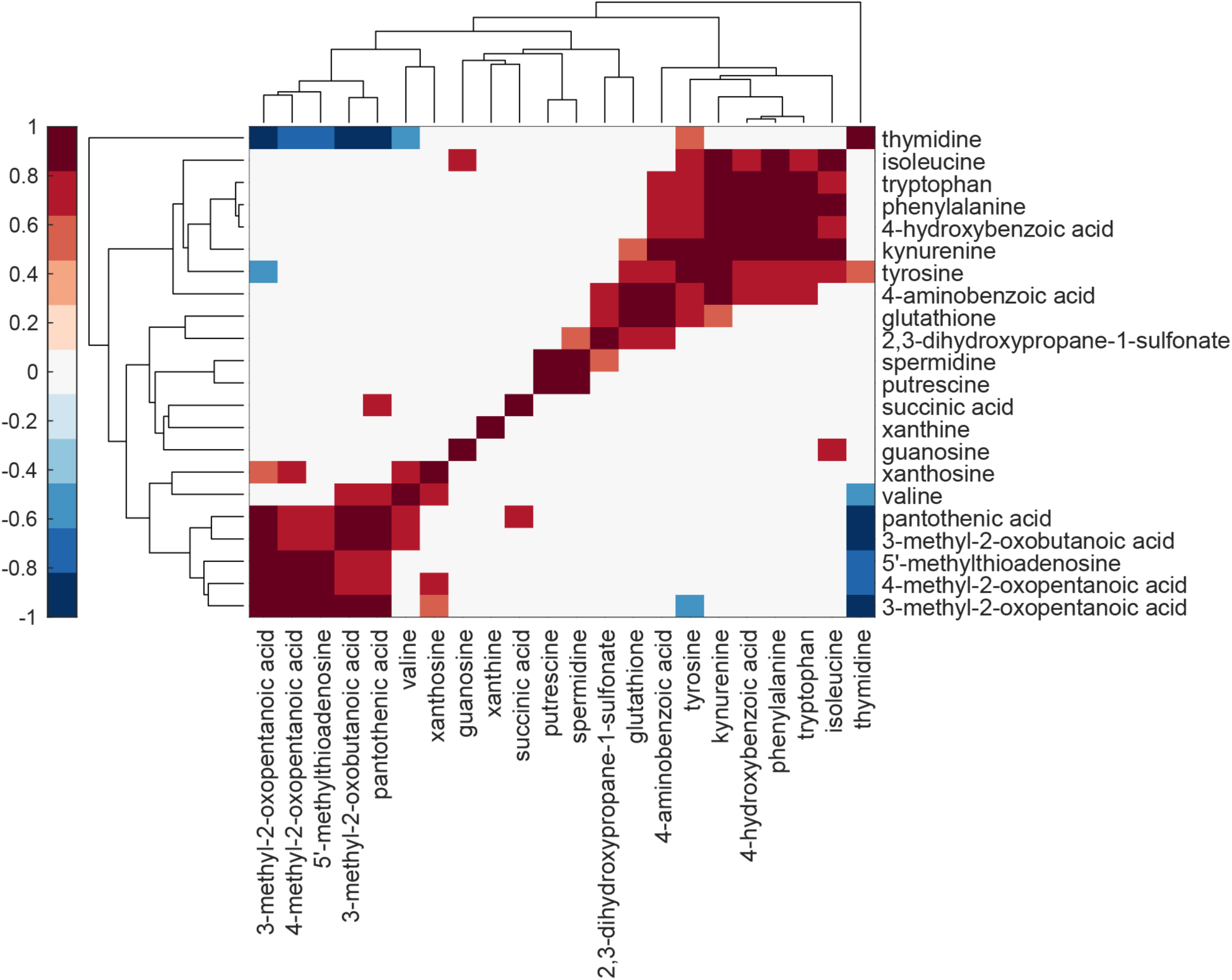
Clustergram of correlations for all dissolved metabolites collected from cells grown in replete conditions. Statistically significant positive (red) and negative (blue) correlations are Pearson correlations with p-values adjusted using a False Discovery Rate of 5%.

**Table S1.** Complete set of metabolites within the targeted metabolomics method used in the current project and the extraction efficiency information from Johnson et al. (2017) which have been updated with unpublished data. Metabolites marked with ‘yes’ in intracellular and/extracellular columns were found in at least one *Prochlorococcus* strain under any light condition. We report all detected metabolites, but only concentrations for those with extraction efficiencies above 1%. Intracellular concentrations of oxidized glutathione are not available due to interference by an unknown compound. The concentration data for each metabolite is available at MetaboLights (http://www.ebi.ac.uk/metabolights/) as study accession number MTBLS567.

| metabolite | extraction efficiency (%) | intracellular | extracellular |
| --- | --- | --- | --- |
| 2,3-dihydroxybenzoic acid | 100.9 |  |  |
| 2,3-dihydroxypropane-1-sulfonate | 0.6 |  | yes |
| 3-mercaptopropionic acid | 88.6 |  |  |
| 3-methyl-2-oxobutanoic acid | 10.1 |  | yes |
| 3-methyl-2-oxopentanoic acid | 50.1 | yes | yes |
| 4-aminobenzoic acid | 18.5 | yes | yes |
| 4-hydroxybenzoic acid | 88 |  | yes |
| 4-methyl-2-oxopentanoic acid | 43.4 | yes | yes |
| 5'-methylthioadenosine | 80.9 | yes | yes |
| 4-amino-5-aminomethyl-2-methylpyrimidine | 0 |  |  |
| D-glucosamine 6-phosphate | 0 |  |  |
| dimethylsulfoniopropionate | 0 |  |  |
| γ-aminobutyric acid | 0 | | |
| 4-Amino-2-methyl-5-pyrimidinemethanol | 0 |  |  |
| nicotinamide adenine dinucleotide | 20.9 | yes |  |
| putrescine | 0 |  | yes |
| acetyltaurine | 0 |  |  |
| adenine | 0 |  |  |
| adenosine | 6.5 |  |  |
| adenosine 5'-monophosphate | 0.2 | yes |  |
| alpha-ketoglutaric acid | 0 | yes |  |
| arginine | 0 | yes |  |
| aspartic acid | 0 | yes |  |
| glycine betaine | 0 | yes |  |
| biotin | 52.8 |  |  |
| caffeine | 23.5 |  |  |
| chitobiose | 0.3 | yes |  |
| chitotriose | 5.6 |  |  |
| choline | 0 |  |  |
| ciliatine | 0 |  |  |
| citrate | 0.9 |  |  |
| citrulline | 0 | yes |  |
| cyanocobalamin | 79.1 |  |  |
| cysteine | 0 |  |  |
| cytosine | 0 |  |  |
| desthiobiotin | 6.5 |  |  |
| dihydroxyacetone phosphate | 0 | yes |  |
| ectoine | 0 |  |  |
| folic acid | 41.2 |  |  |
| fosfomycin | 0 |  |  |
| fumaric acid | 0 |  |  |
| glucose 6-phosphate | 0 | yes |  |
| glutamic acid | 0 | yes |  |
| glutamine | 0 | yes |  |
| glyphosate | 15.1 |  |  |
| glutathione | 1.1 | yes | yes |
| glutathione oxidized | 1.5 | not available | yes |
| guanine | 0 |  |  |
| guanosine | 7.7 | yes | yes |
| hemin | 0 | yes |  |
| indole 3-acetic acid | 16.8 |  |  |
| inosine | 8.1 |  |  |
| inosine 5'-monophosphate | 0 | yes |  |
| isoleucine | 1.41 | yes | yes |
| kynurenine | 41.7 |  | yes |
| leucine | 3.3 | yes |  |
| malic acid | 0.7 |  |  |
| methionine | 0 | yes |  |
| muramic acid | 0 |  |  |
| n-acetyl glucosamine | 0 |  |  |
| n-acetyl glutamic acid | 1.1 | yes |  |
| n-acetyl muramic acid | 2.8 |  |  |
| ornithine | 0 |  |  |
| orotic acid | 0 |  |  |
| pantothenic acid | 51.9 | yes | yes |
| phenylalanine | 39.7 | yes | yes |
| phosphoenolpyruvate | 0 |  |  |
| phycocyanobilin | 0 |  |  |
| proline | 0 | yes |  |
| pyridoxine | 6.8 |  |  |
| riboflavin | 87.6 |  |  |
| S-(5'-adenosyl)-L-homocysteine | 43.3 | yes |  |
| S-adenosyl-L-methionine | 0 | yes |  |
| alanine (isom. sarcosine) | 0 |  |  |
| serine | 0 |  |  |
| sn-glycerol 3-phosphate | 0 |  |  |
| taurocholic acid | 92.8 |  |  |
| spermidine | 0 | yes | yes |
| succinic acid | 0 |  | yes |
| syringic acid | 32.9 |  |  |
| taurine | 0 |  |  |
| thiamine | 2 |  |  |
| thiamine monophosphate | 0 |  |  |
| threonine / homoserine | 0 |  |  |
| thymidine | 52.6 |  | yes |
| tryptamine | 21.3 |  |  |
| tryptophan | 46.7 | yes | yes |
| tyrosine | 2.1 | yes | yes |
| uracil | 0 | yes |  |
| uridine 5'-monophosphate | 0 | yes |  |
| valine | 0 | yes | yes |
| xanthine | 0.3 |  | yes |
| xanthosine | 10.4 |  | yes |

**Table S2.** Mean (± one standard deviation) cell-specific concentrations of glycine betaine (fg cell1) in cultures of three strains of *Prochlorococcus* grown at a range of light intensities. *Biomass of cells from each strain are from Cermak et al. (2017). We assumed 50% of the cell was carbon. These values were used to calculate the percent of each strain’s intracellular carbon content that can be attributed to the carbon in glycine betaine. †Measured value for NATL2A, a related LLI strain of *Prochlorococcus*.

| Strain | Clade | Growth light intensity ($\mu\text{mol photons m}^{-2}\text{s}^{-1}$) | Median cellular biomass ($\text{fg}$)* | Glycine betaine ($\text{fg cell}^{-1}$) | % of biomass |
| --- | --- | --- | --- | --- | --- |
| MIT9301 | HLII | 10 | $60 \pm 3$ | $4.5 \times 10^{-4} (\pm 8.0 \times 10^{-4})$ | 0.002% |
| MIT9301 | HLII | 50 | $60 \pm 3$ | 0 | |
| MIT0801 | LLI | 10 | $91 \pm 5^{\dagger}$ | 0 | |
| MIT9313 | LLIV | 5 | $158 \pm 6$ | $3.0 (\pm 1.3)$ | 4% |
| MIT9313 | LLIV | 10 | $158 \pm 6$ | $4.7 (\pm 2.4)$ | 6% |

